# Ribosome reinitiation can explain length-dependent translation of messenger RNA

**DOI:** 10.1101/103689

**Authors:** David W. Rogers, Marvin A. Böettcher, Arne Traulsen, Duncan Greig

## Abstract

Models of mRNA translation usually presume that transcripts are linear; upon reaching the end of a transcript each terminating ribosome returns to the cytoplasmic pool before initiating anew on a different transcript. A consequence of linear models is that faster translation of a given mRNA is unlikely to generate more of the encoded protein, particularly at low ribosome availability. Recent evidence indicates that eukaryotic mRNAs are circularized, potentially allowing terminating ribosomes to preferentially reinitiate on the same transcript. Here we model the effect of ribosome reinitiation on translation and show that, at high levels of reinitiation, protein synthesis rates are dominated by the time required to translate a given transcript. Our model provides a simple mechanistic explanation for many previously enigmatic features of eukaryotic translation, including the negative correlation of both ribosome densities and protein abundance on transcript length, the importance of codon usage in determining protein synthesis rates, and the negative correlation between transcript length and both codon adaptation and 5' mRNA folding energies. In contrast to linear models where translation is largely limited by initiation rates, our model reveals that all three stages of translation - initiation, elongation, and termination/reinitiation - determine protein synthesis rates even at low ribosome availability.

## Introduction

The physiological state of a cell is largely determined by the identity and abundance of the proteins encoded by its genome. Understanding how genetic information is first transcribed into messenger RNA and then translated into protein is therefore fundamental to our understanding of biological systems. A wide variety of technologies has allowed detailed investigations of transcription, but - until very recently - a lack of similar tools for empirical research on translation has meant that the study of post-transcriptional regulation has been largely restricted to mathematical models with little opportunity for parameterization or evaluation. Recent advances in both sequencing technology and mass spectrometry have now produced large amounts of data on the translation of eukaryotic mRNA, revealing how transcript features, RNA-binding proteins, and non-coding RNAs influence translation [1,2]. While many of the determinants of translation rates revealed by these empirical studies were predicted by existing models, some remain difficult to explain. Perhaps the most striking correlate of translation rate is the length of the transcript itself. Multiple experimental studies, across a wide range of eukaryotic organisms, have demonstrated a steep negative correlation between the length of a given coding sequence (CDS) and three different measures of translation: translation initiation rates [3#5], the density of ribosomes on a transcript [5–15], and the abundance of the encoded protein [16–19].

Ribosome and polysome profiling experiments have shown a positive relationship between ribosome density and protein abundance, leading to the conclusion that transcripts with higher ribosome densities have higher translation rates [9,11,20]. A positive relationship between ribosome density and translation rate can occur when translation is limited by low initiation rates. In traditional models of translation, initiation can be limiting when other steps in translation, such as elongation, occur quickly enough to prevent collisions between ribosomes [20]. Consistent with this key role of initiation rates in determining translation rates, Arava et al [6] found that the higher densities of ribosomes on shorter transcripts was most consistent with shorter transcripts having exponentially higher initiation rates than longer transcripts, estimating a halving of the initiation rate with every 400-codon-increase in CDS length. More recent analyses [3,4] have revealed that the relationship between CDS length and initiation rates is better described by a power law: the initiation rate is roughly halved for every doubling of CDS length (i.e. a log-log slope of approximately −1). However, the assumption of initiation-limitation leaves little room for variation in elongation rates to influence translation rates, which is at odds with recent work demonstrating that codon usage can be an important determinant of protein yields [21,22].

If translation is limited by the ability of transcripts to capture ribosomes from the cytoplasmic pool (the *de novo* initiation rate), mechanisms that allow transcripts to retain terminating ribosomes for subsequent rounds of translation should improve translation rates. The closed-loop model of translation was first proposed as a hypothetical mechanism to improve translation efficiency through intrapolysomal ribosome reinitiation [23,24]. By bringing the sites of termination and initiation into close proximity through circularization of the mRNA, the closed-loop complex allows ribosomes that have finished translating to reinitiate translation on the same mRNA molecule rather than returning to the cytoplasmic pool. The closed-loop model was initially based on the appearance of many polysomes in electron micrographs as circular, rather than linear, structures (detailed high resolution tomographic analyses of circular polysomes are now available [25]). Recent theoretical and experimental studies have shown that secondary structures in single stranded RNAs bring the 5) and 3) ends close together (equivalent to the distance spanned by 9-16 nucleotides) meaning that mRNAs are effectively circularized [26,27]. Interactions between initiation factors bound to the 5' end, and proteins associated with the 3' end including release and recycling factors, and the poly(A) binding proteins, are thought to facilitate translation, possibly by stabilizing the closed-loop structure or by actively promoting reinitiation [28,29].

The importance of reinitiation of ribosomes on circular transcripts in determining protein yield is well established *in vitro* [23,24,30–32]. Measuring translation of the luminescent protein luciferase in a eukaryotic cell-free system, Kopeina et al [31] showed that circular polysomes rarely exchanged ribosomes with the free pool or lost ribosomes to other transcripts, but linear polysomes did so frequently. On circular polysomes, most terminating ribosomes immediately reinitiated on the same mRNA molecule (see also [30]). Alekhina et al [32] found that protein production in a similar cell-free system does not rapidly reach a steady state, as would be expected under a linear model of translation, but rather accelerates over the lifetime of the transcript, consistent with reinitiation on the same transcript. They proposed that the translation rate initially depends on slow *de novo* initiation of ribosomes from the free pool but soon becomes dominated by the much faster process of reinitiation.

Here, we use a minimal computational model to investigate the consequences of ribosome reinitiation on translation, with particular focus on transcript length and codon usage. We find that reinitiation causes ribosome densities, overall initiation rates, and protein yields to decrease with increasing transcript length. Furthermore, higher levels of reinitiation increase the importance of codon usage in determining translation rates in a length-dependent manner, even at low ribosome densities or low *de novo* initiation rates. Reinitiation therefore provides a potential mechanistic explanation for multiple previously-enigmatic patterns observed in empirical studies of translation.

## Model Description

We use a totally asymmetric simple exclusion process (TASEP, reviewed in [33]) to investigate the closed loop model of translation. The TASEP (Fig. 1) models each transcript as a one-dimensional lattice consisting of a number of sites equal to the number of codons in the CDS: each site represents a single codon. Each site can be either free or occupied by a ribosome. Ribosomes move along the transcript in the 5′ to 3′ direction and cannot occupy the same codon(s) as any other ribosome. In our model, the transcript is circularized, meaning that terminating ribosomes can not only be released into the cytoplasmic pool (as in a linear TASEP) but can also move to the initiation site of the same transcript (reinitiation).

**Figure 1.**
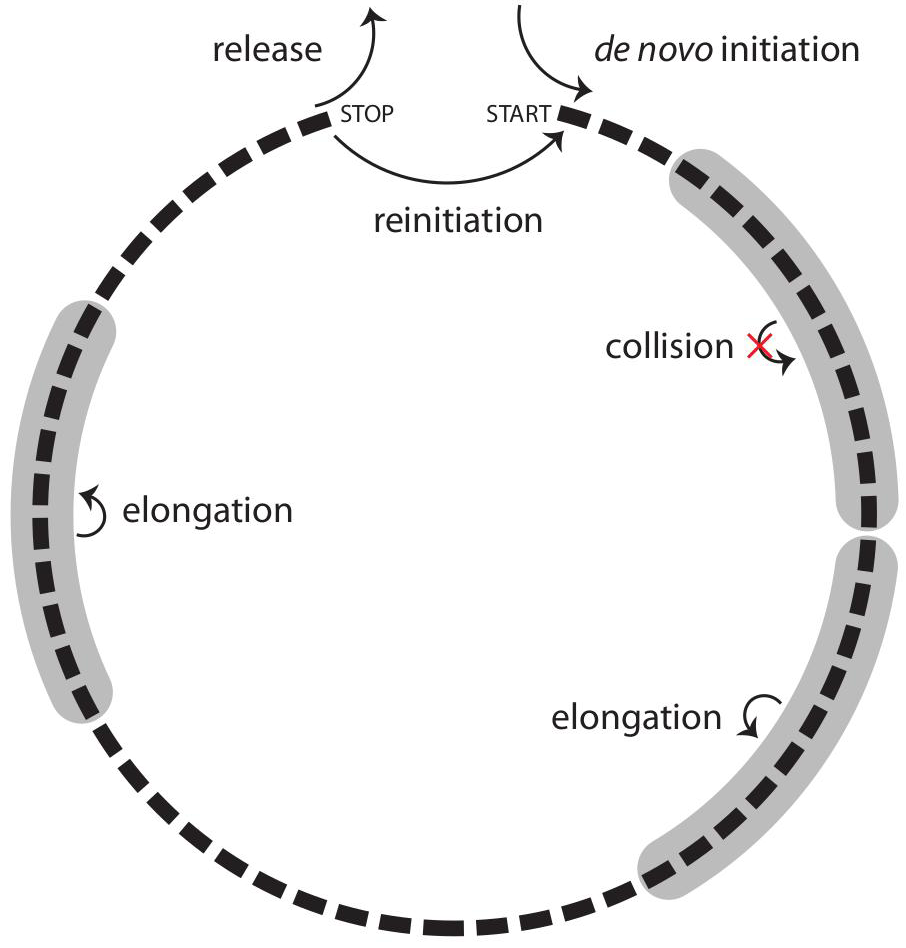
The closed-loop model of translation. Ribosomes (shaded in grey), modeled as extended particles that occupy 10 codons (black boxes), join the transcript at the start codon from the cytoplasmic pool at the *denovo* initiation rate, and hop to the next codon (in the 5' to 3' direction) at the elongation rate. Upon reaching the stop codon, ribosomes either return to the cytoplasmic pool at the release rate or return to the start codon at the reinitiation rate. The reinitiation level is determined by the reinitiation rate divided by the sum of the reinitiation and release rates. If the initiation site is occupied (i.e. any of the first 10 codons is being decoded), new ribosomes fail to join the transcript and reinitiating ribosomes either remain at the termination site or return to the cytoplasmic pool at the release rate. An elongating ribosome fails to step forward if the distance between its center and that of the ribosome in front is ≤ 10 codons (collision). Stochastic simulations were performed using the Gillespie algorithm. The Gillespie algorithm consists of multiple steps: (1) Initialization: the simulation time is set to zero; at this point in our simulations, all transcripts are empty and the only possible reaction is *de novo* initiation; (2) List all possible reactions: all possible reactions are determined and their rates are used to calculate the total rate of possible reactions; (3) Monte Carlo step: two random numbers are generated, the first determines the waiting time until the next reaction based on the total rate of possible reactions, and the second determines which reaction occurs using each reaction rate as a probabilistic weight; (4) Update: the time is increased by the randomly generated waiting time from step 3 and the chosen reaction is performed; (5) Iteration: repeat from step 2 unless the transcript lifetime has been reached.

Four different types of reactions can take place in the TASEP: (i) *de novo* initiation: a free ribosome can be placed onto the 5' end of the transcript (the initiation site) at the *de novo* initiation rate; (ii) elongation: ribosomes at any codon on the transcript (except the termination site) can move forward one codon in the 3' direction at the elongation rate; if a ribosome occupies the termination site, it can either (iii) leave the transcript at the release rate or (iv) it can move to the initiation site at the reinitiation rate.

We model ribosomes as extended particles that occupy ten codons each: the A-sites (where each codon is translated) of adjacent ribosomes must be spaced apart by at least 10 codons. Thus, the elongation reaction is only possible when the A-site of the next ribosome in the 3' direction is > 10 codons downstream. Similarly, neither *de novo* initiation nor reinitiation is possible if any of the first 10 codons is occupied by an A-site.

Analytical solutions of the TASEP are possible, but currently can only be applied to the steady state. Consequently, most TASEP models, including a recent study of reinitiation [34], investigate translation at the steady state, where the rate at which ribosomes join a transcript equals the rate at which they leave, and the translation rate is constant. However, every real transcript spends some proportion of its lifetime outside of the steady state, where these solutions do not apply; the assumption of a perpetual steady state is therefore an approximation. A new transcript does not instantly acquire ribosomes distributed along its length. Instead, ribosomes join at the 5′ end and gradually progress towards the stop codon, where they can be released. In the absence of reinitiation, the steady state can be reached once the first ribosome to join a transcript is released. The duration of this ,pioneer round- increases with transcript length, but generally represents a small proportion of eukaryotic transcript lifetimes. In the absence of reinitiation the steady state is therefore a good approximation (although it can be inappropriate for prokaryotes with short-lived transcripts [36,37]). However under reinitiation, ribosomes do not necessarily leave the transcript upon termination, which causes the effective initiation rate (and the translation rate) to increase over time [32]. The time to reach the steady state therefore increases with both transcript length and reinitiation probability, and the time spent outside of the steady state thus represents a greater proportion of transcript lifetimes (Fig. S1A). The steady state assumption consequently becomes a much worse approximation of translation at higher levels of reinitiation, overestimating translation rates on long transcripts and underestimating translation on short transcripts (Fig. S1B). It is therefore impossible to make a fair comparison of translation at different reinitiation levels using the steady state approximation, particularly for transcripts of different lengths.

Since we do not assume that translation on any given transcript is always at the steady state, we cannot use the steady state analytical solutions of the TASEP. Instead, we perform stochastic simulations using the Gillespie algorithm [38], which capture both the steady state and the non-steady state. In models that assume the steady state, all translation that occurs in simulations prior to the steady state is ignored. For example, in a recent reinitiation-based model of translation in yeast, the first 10^5^ s of simulations was discarded [34]. Given that the average lifetime of yeast transcripts is on the scale of 10^3^-10^4^ s [39], this means that all translation occurring over biologically plausible lifetimes was excluded from the analysis. Here, we make no assumptions about the steady state; we simply account for all translation that occurs during the lifetime of a transcript (both before and after the steady state is achieved). We simulated translation on each transcript independently. Each run generated a time evolution of the ribosome occupancy at each codon on a given transcript. We computed three measures of translation: ribosome density (the average number of ribosomes on a transcript over its lifetime divided by one tenth of CDS length, because each ribosome occupies 10 codons), effective initiation rate (the total number of initiations occurring through either *de novo* initiation or reinitiation divided by transcript lifetime) and protein yield (the total number of ribosomes reaching the stop codon of a transcript). We averaged the results of 1000 runs to produce results that are not subject to large stochastic fluctuations. We do not consider untranslated regions and our transcripts therefore represent only the CDS. The code for our TASEP is available at: https://github.com/marvinboe/reTASEP

Changing any transcriptome-wide parameter can dramatically alter global ribosome usage. For instance, at a given *de novo* initiation rate, increasing the reinitiation probability increases the total number of actively translating ribosomes. While this effect may be true, given that reinitiation is expected to allow more efficient use of ribosomes (see Discussion), it makes parameterizing the model difficult because the actual level of reinitiation is unknown. To keep all simulations consistent with empirical values, we have adjusted the *de novo* initiation rate to maintain the empirically observed average ribosome density. For simplicity, we have kept the number of ribosomes on a 400-codon-long transcript constant (at 6 ribosomes) for all transcriptome-wide reinitiation probabilities and elongation rates.

## Results

### High levels of reinitiation generate length-dependent translation

Our model captures the negative correlation between ribosome density and CDS length observed in empirical studies, but only if the probability of reinitiation is high (Fig. 2). This result is intuitive; if reinitiation were perfect, all ribosomes that initiate would continue to reinitiate and translate, never leaving a transcript until it degrades. The density of ribosomes on a transcript of a given length and age would therefore be determined exclusively by the *denovo* initiation rate. If the *de novo* initiation rate is the same for all transcripts, then all transcripts of a given age should carry the same number of ribosomes and ribosome density will be the inverse of CDS length (with a log-log slope of −1). At a given elongation rate, the time required for a ribosome to complete one cycle (travel from the start codon to the stop codon) is less for short transcripts than for long transcripts. This means that, prior to the steady state, reinitiation occurs more frequently on shorter transcripts resulting in higher protein yields for short transcripts than long transcripts. When all or nearly all terminating ribosomes reinitiate, the effective initiation rate is much higher for shorter transcripts - providing a simple mechanism that could explain the length-dependence of initiation rates predicted by recent studies of translation [3–5,8]. The higher ribosome densities on shorter transcripts, and corresponding higher levels of reinitiation, result in higher occupancies of the initiation site on shorter transcripts, occasionally blocking new ribosomes from initiating on short transcripts. This initiation interference [21] slightly reduces the length-dependence ofribosome density, effective initiation rate, and protein yield (Fig. 2), resulting in a log-log slope less steep than −1.

When reinitiation is not perfect, ribosomes can return to the cytoplasmic pool after termination, and the effect of CDS length on ribosome density, effective initiation rate, and protein yield is diminished. Even small reductions in reinitiation probability greatly weaken length-dependence (Fig. 2). This is because short transcripts have more opportunities to lose ribosomes than do long transcripts. While a successful reinitiation event only guarantees that a ribosome remains associated with the transcript until the next termination event, ribosome loss is permanent. In the complete absence of reinitiation, length-dependence is therefore abolished.

**Figure 2.**
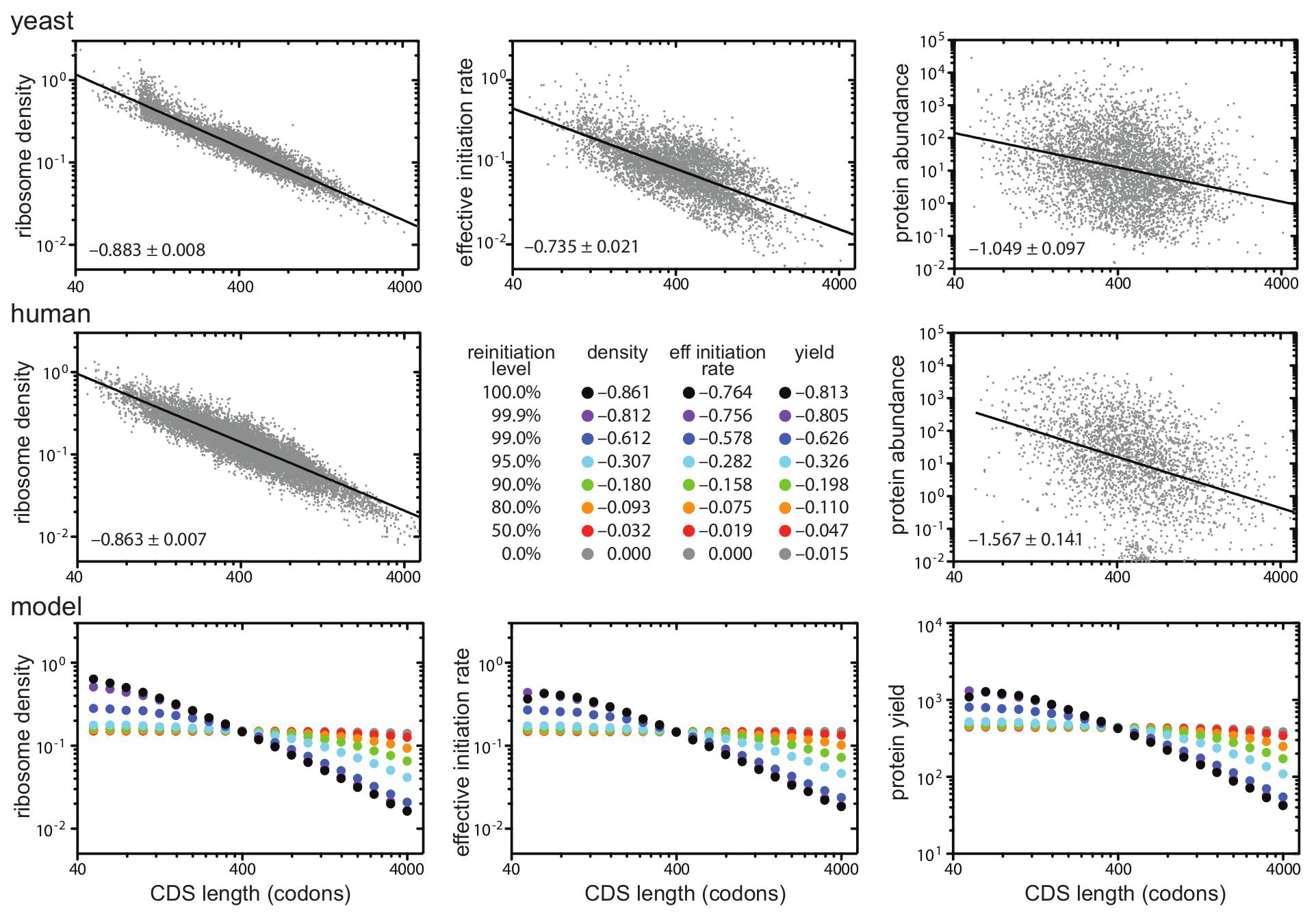
Reinitiation of post-termination ribosomes causes length-dependent translation. Ribosome density is the average number of ribosomes occupying a transcript during its lifetime divided by one tenth of the CDS length (since each ribosome occupies 10 codons). The effective initiation rate is the total number of initiation events (*de novo* initiation and reinitiation) divided by transcript lifetime. Protein yield is the total number of ribosomes reaching the stop codon during the lifetime of a transcript. Slopes (95% confidence intervals) are indicated in the bottom left corner of each panel. *De novo* initiation rates were adjusted at each reinitiation level so that a 400-codon long transcript carried 6 ribosomes. **Top**: experimentally observed relationships between CDS length and ribosome density (left), initiation rate (center) or protein abundance (right) in the budding yeast *Saccharomyces cerevisiae*.**Middle**: experimentally observed relationship between CDS length and ribosomedensity (left) and protein abundance in the human HEK293T cell line. Estimates of the initiation rate are not currently available for this cell line, so we have used this space to list the slopes from our simulations. **Bottom:** predicted relationships between ribosome density (left), the effective initiation rate (center), and protein yield (right) and CDS length at different reinitiation levels (different colours) from our simulations. Simulations were performed using a fixed elongation rate of 10s^-1^ (see Fig. S3 for simulations at other fixed elongation rates). **Datasources**: Yeast densities are weighted averages of the signals in polysomal fractions for 6071 transcripts from[6]; initiation rates for 5348 transcripts were calculated by Ciandrini et al [3] based on ribosome density data from [7]; protein abundances of 4694 proteins included in the Peptide Atlas 2013 dataset from PaxDb [40] normalizedagainst the total number of proteins (expressed as parts per million). HEK293T densities were calculated from mean ribosome numbers (across 3 replicates) reported by Hendrickson et al [11]; protein abundances of 2636 proteins with identified CDS lengths included in the Geiger MCP 2012 data set (based on spectral counting) from PaxDb [40].

### Reinitiation, but not *de novo* initiation, has a larger effect on short transcripts than long transcripts

While changing transcriptome-wide parameters can dramatically affect global ribosome usage (see Model Description), altering parameters of transcripts encoded by a single gene will have little effect on global ribosome usage. This is because nearly all endogenous genes are expressed at low levels, so changing the translation parameters of the transcripts produced by a single gene will have a negligible effect on global ribosome availability [4,20,41]. By studying transcripts of individual genes, we can therefore investigate the consequences of changing a single parameter while holding all other values constant. We first tested the effects of altering the reinitiation rate of transcripts encoded by a single gene (Fig. 3 A, B). Doubling the reinitiation rate results in an extremely similar increase in all three measures of translation (ribosome density, effective initiation rate, and protein yield; results are therefore only shown for ribosome density), but the effects are greater for short transcripts than long transcripts. These effects are mirrored by a length-dependent decrease in translation when the reinitiation rate is halved (Fig. 3B). Furthermore, the length-dependent effects of changing the reinitiation rate of a single transcript species are generally stronger at higher transcriptome-wide reinitiation probabilities, except when reinitiation is so high that ribosomes rarely leave the transcript (e.g. 99.9%).

We next tested the effects of altering the *de novo* initiation rate of a single transcript species (Fig 3 C, D). In the absence of reinitiation, doubling the *de novo* initiation rate had an equal effect on ribosome density for transcripts of all lengths. However, at higher levels of reinitiation, doubling the *de novo* initiation rate resulted in a smaller increase in ribosome density on short transcripts than on long transcripts, caused by increased initiation interference; the higher density of ribosomes on short transcripts under reinitiation increases the probability that the initiation site is blocked, preventing successful *de novo* initiation. The effects of altering the *de novo* initiation rate on the effective initiation rate and protein yield are very similar to the effects on ribosome density.

**Figure 3.**
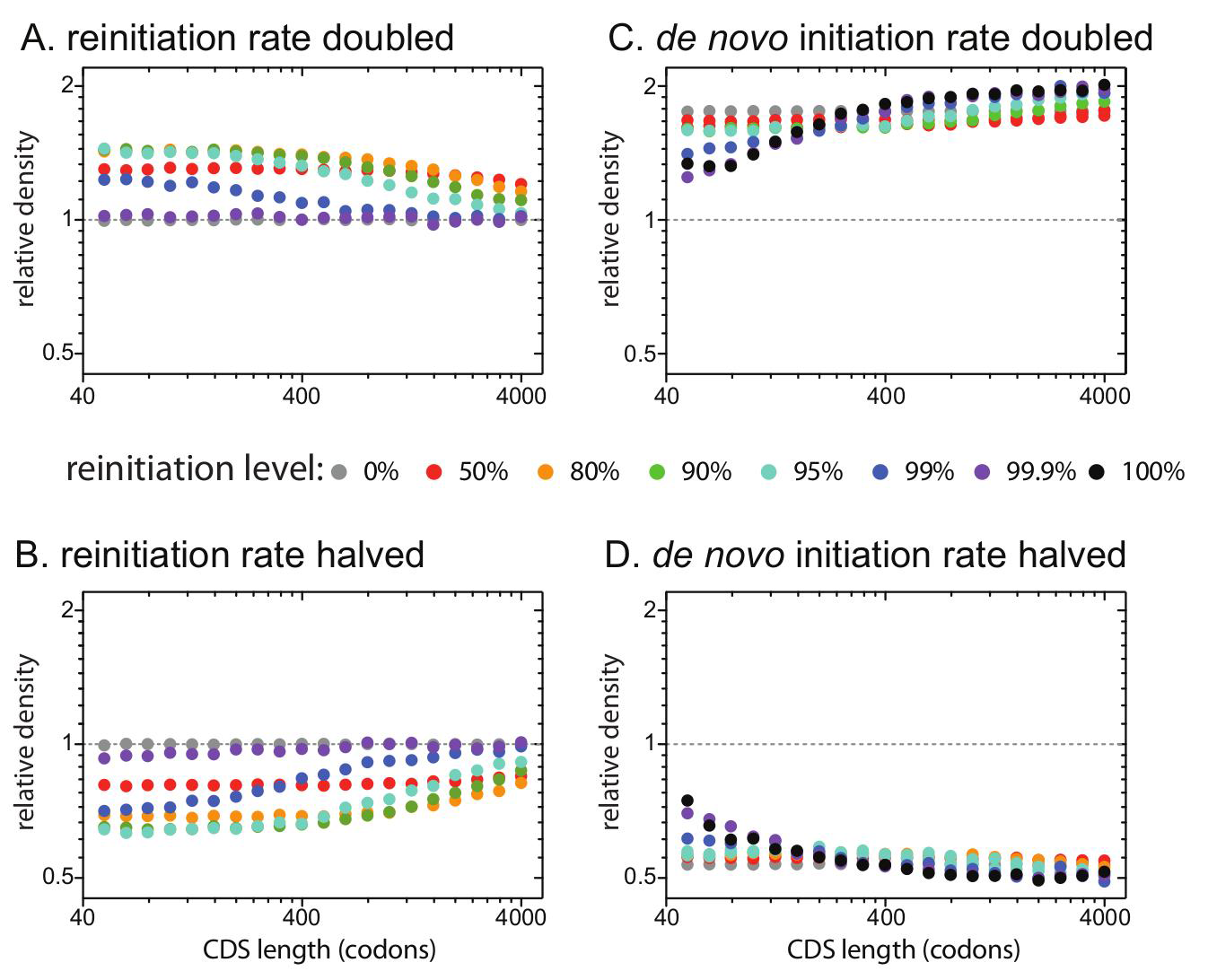
Transcript-specific change of the reinitiation rate, but not the *de novo* initiation rate, has larger effects on short transcripts than long transcripts. We simulated the effects of changing the reinitiation rate (A,B or the *de novo* initiation rate (C, D) of a single transcript species by2-fold or0.5-fold at different transcriptome-wide reinitiation levels. For each transcriptome-wide reinitiation level (different colours), doubling (or halving) the reinitiation rate shifted the reinitiation level to:99% to99.5% (98.0%);95% to97.5% (90.5%);90% to94.7% (81.8%);80% to88.9% (66.7%);50% to66.7% (33.3%);0% to 0% (0%). Doubling or halving the reinitiation rate at very high transcriptome-wide reinitiation levels (e.g. 99.9%) has little effect on translation since ribosomes rarely leave transcripts. Y-axes (log2 scaled) show the ribosome density of altered transcripts relative to an equivalent transcript at the transcriptome wide reinitiation level. The effects of changing either the reinitiation rate or the *de novo* initiation rate on the effective initiation rate and protein yield were nearly identical to the effects on ribosome density. Transcriptome-wide *de novo* initiation rates were adjusted at each reinitiation level so that a 400-codon long CDS at the transcriptome-wide reinitiation level carried6 ribosomes.

### High levels of reinitiation couple effective initiation rates and protein yields to the elongation rate

So far, we have assumed that all transcripts have identical elongation rates, but in reality the elongation rate varies between transcripts encoded by different genes [42]. We therefore investigated the consequences of changing the elongation rate of a single CDS from 10s^-1^ to either 20s^-1^ or 5s^-1^ (Fig. 4). Increasing the elongation rate reduces the amount of time between initiation and termination. In the absence of reinitiation, this causes ribosomes to spend less time on the altered transcript resulting in decreased ribosome density, but has little effect on the initiation rate or protein yield since these elongation rates are generally not limiting. Altered elongation rates do affect how long it takes to clear the initiation site and therefore the amount of initiation interference, explaining the relatively small differences in initiation rates and protein yields seen at 0% reinitiation [21].

Under perfect reinitiation, terminating ribosomes explicitly reinitiate on the same transcript. Changing the elongation rate of a single gene therefore has no effect on the density of ribosomes on the altered transcript. However, by altering the time between reinitiation events, changing the elongation rate results in an equal change in the effective initiation rate of the altered transcript (Fig. 4). The protein yield of any endogenous gene is therefore exquisitely sensitive to changes in elongation rate under perfect reinitiation. Under perfect reinitiation, this effect is seen at all CDS lengths. The importance of the elongation rate decreases dramatically when reinitiation levels are reduced: faster elongation results in more opportunities to lose ribosomes, particularly on short transcripts.

**Figure 4.**
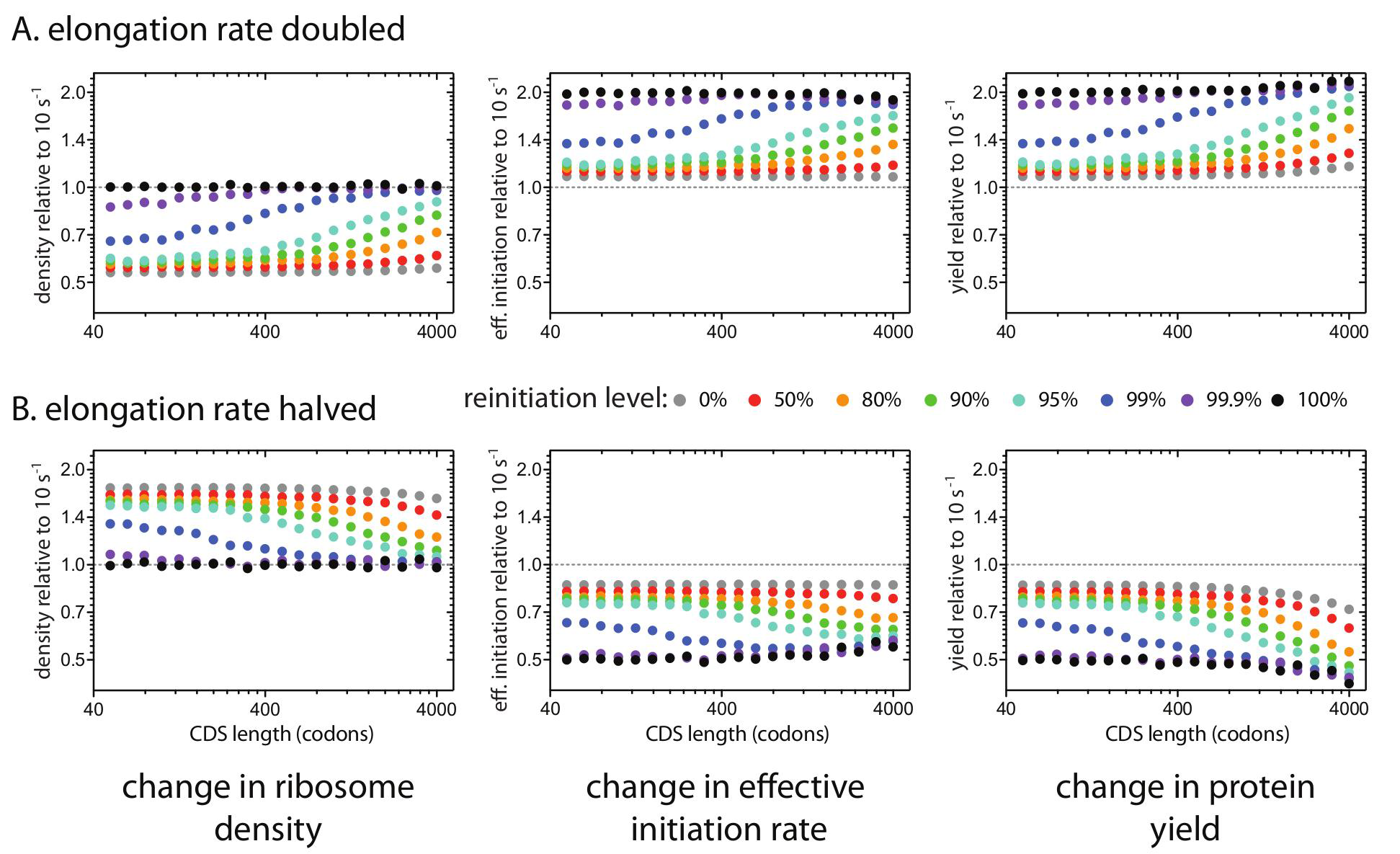
Transcript-specific change in translation caused by altering the average elongation rate of a single coding sequence. We simulated the effects of changing the average elongation rate of a single transcript species from 10s^-1^ to either (A) 20s^-1^ or (B) 5s^-1^ at different reinitiation levels (different colours). CDS length refers to the length of the altered coding sequence. Y-axes (log 2 scaled) show the effect of altering the elongation rate on the ribosome density (left), effective initiation rate (center), and protein yield (left) of the altered transcript at the higher or lower elongation rate relative to 10s^-1^ (dotted line). *De novo* initiation rates were adjusted at each reinitiation level so that a 400-codon long transcript with a fixed elongation rate of 10s^-1^ carried 6 ribosomes.

### Length-dependent consequences of a single slow step on translation

So far, we have only considered the effects of changing the average elongation rate of a transcript. However, it is difficult to imagine a mechanism that could simultaneously alter the elongation rate of all codons in a single transcript species without affecting the global elongation rate. Instead, transcripts are likely altered by mutations affecting a single codon at a time. Codon usage can affect elongation by determining the stability of secondary structures in the mRNA, but different codons are also decoded at different rates depending on the cellular availability of the appropriate tRNA. Most amino acids are encoded by multiple codons, and some codons (including synonymous codons that code for the same amino acid) are decoded faster than others [42,43]. We therefore investigated the consequences of a single slow step on translation of transcript species of different lengths (Fig. 5). Here, we only examined translation at 99.9% reinitiation; similar results would be expected for other models of length-dependent translation. Introducing a single slow step into any transcript reduces its effective initiation rate and protein yield, but the effects are much larger for short transcripts than for long transcripts (Fig. 5). The length-dependence of a single slow step arises from two sources. First, a single site represents a larger proportion of a short transcript than a long transcript and consequently results in a greater decrease in the average elongation rate [46]. Second, short transcripts have higher ribosome densities and are therefore more prone to collisions or "traffic jams" than are long transcripts. Effective initiation rates and protein yields are particularly sensitive to single slow steps near the start codon, with larger effects on shorter transcripts: slow clearance of the initiation site delays reinitiation and blocks *de novo* initiation resulting in lower ribosome densities on affected transcripts.

**Figure 5.**
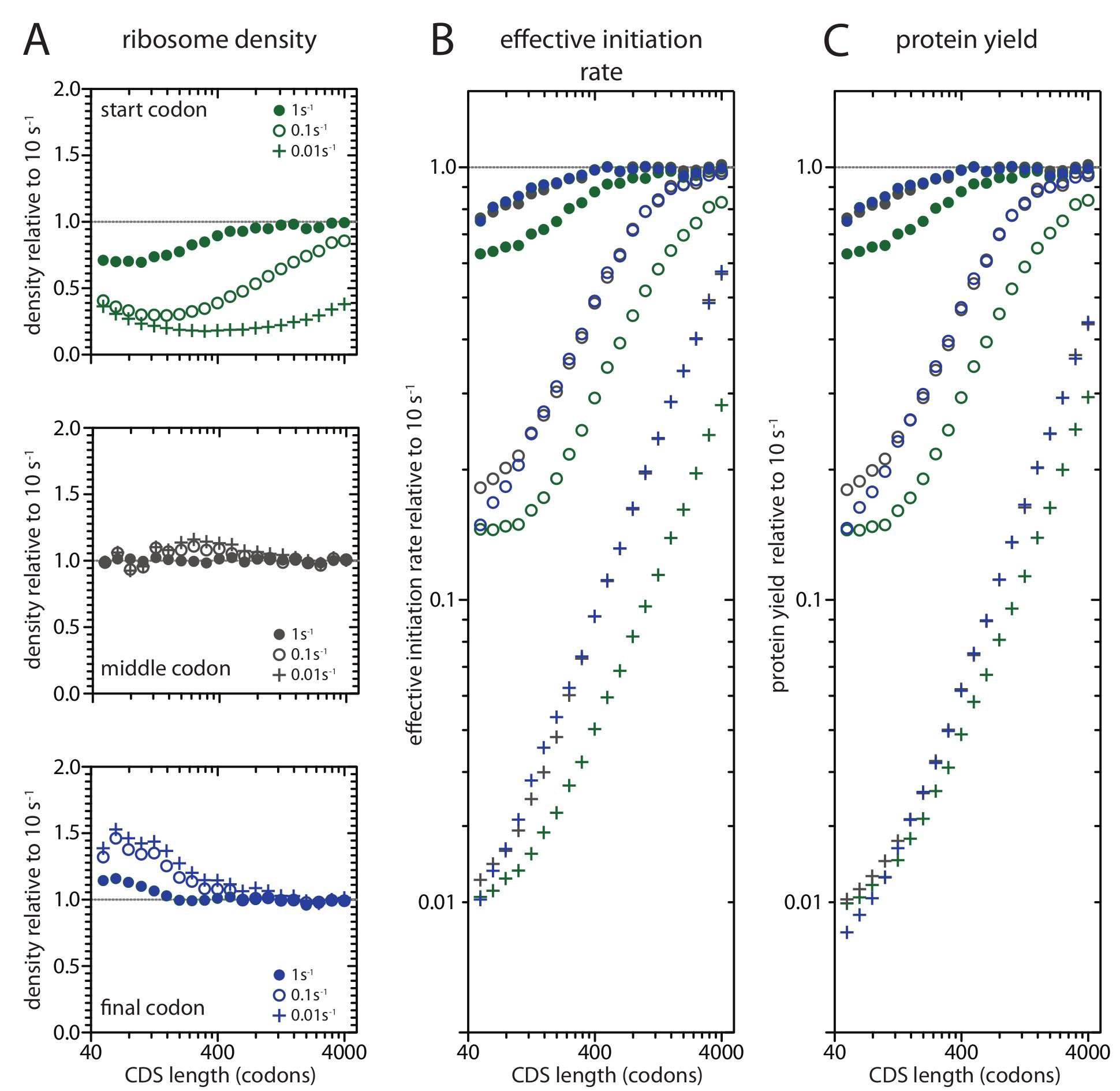
The consequences of a single slow step under length-dependent translation. We investigated the consequences of introducing a slow elongation step at either the start codon (green), the middle codon (grey), or the final codon (immediately before the stop codon, blue) under 99.9% reinitiation. CDS length refers to the length of the altered coding sequence. Slow codons were translated at 1s^-1^ (filled circles), 0.1s^-1^ (open circles), or 0.01s^-1^ (plus signs). Y-axes show the effect of a single slow step on the ribosome density (right), effective initiation rate (center), and protein yield (right) at the lower elongation rate relative to 10s^-1^ (dotted line).

### A yeast-specific model of translation with reinitiation

Given the importance of variation in elongation rates to translation under reinitiation, we used our model to simulate translation in *S. cerevisiae* using codon-specific decoding rates. We used decoding rates (see Table S1) estimated by Gilchrist & Wagner [47] which are based on tRNA availability and wobble pairing rules and scaled so that the average decoding rate is 10s^-1^; they are related to measures of codon occupancy (r = 0.494, n = 61, *P* < 0.0001 [11]). Since efficient reinitiation couples protein production to elongation rates, synonymous codon usage should have detectable consequences for protein yield at high levels of reinitiation. We tested the effects of synonymous codon usage at 99.9% reinitiation by predicting the yields of nine different synthetic GFP constructs [48] that differ only in their synonymous codon usage (Fig. 6A). We compared these predictions to observed protein abundances measured in *S. cerevisiae* expressing each construct, and found a strong positive correlation between predicted yields and observed abundances (r = 0.750, n = 9, *P* = 0.020); our model predicted approximately half of the observed effect of using different synonymous codons (relative expression of highest vs. lowest construct, model = 2.4-fold, observed = 5.4-fold). Thus, efficient reinitiation correctly predicts a role for synonymous codon usage in determining yield.

**Figure 6.**
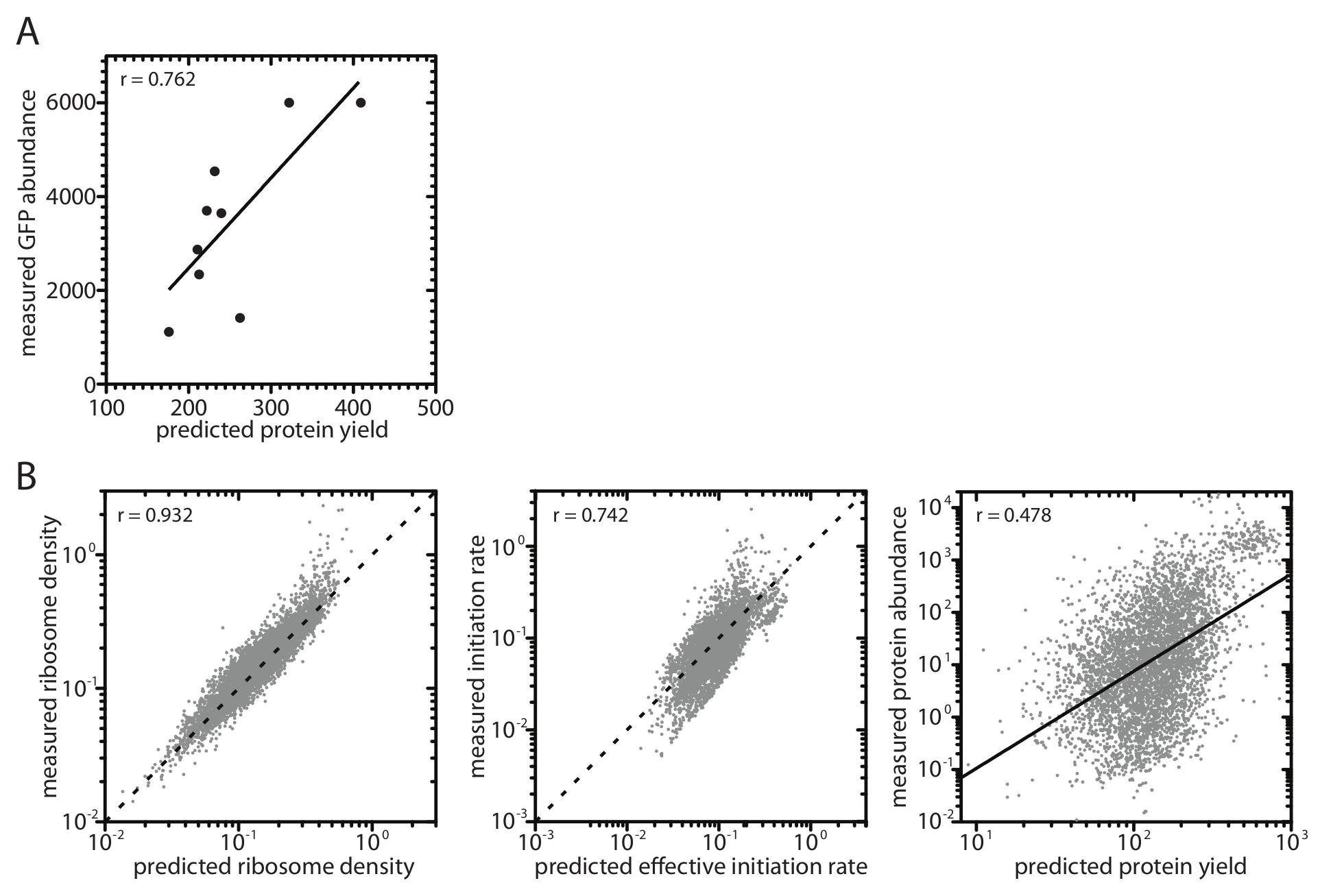
Simulating translation in the budding yeast *S. cerevisiae*. Our simulations were run using yeast-specific decoding rates [47] and a transcript lifetime of 1553s [39]. Results are shown for 99.9% reinitiation (see Figs. S3 & S4 for simulations at other reinitiation levels). (A) Correlation of model-predicted protein yields (proteins per mRNA per lifetime) for 9 differently codon-optimized GFP coding sequences with experimentally measured abundances (arbitrary units) from [48]. (B) Correlation of model-predicted ribosome density, effective initiation rate, and protein yield with experimentally measured values. Sources of experimental data are the same as in Fig. 2. When X and Y-axis scales are equivalent, the 1:1 line is represented by a dotted line. When X and Y-axis scales are not equivalent, the solid line is the regression line.

**Table S1.** .*S. cerevisiae*codon-specific elongation rates from [47].

| Amino acid | Codon | Decoding rate | Amino acid | Codon | Decoding rate | Amino acid | Codon | Decoding rate |
| --- | --- | --- | --- | --- | --- | --- | --- | --- |
| ALA | GCU | 18.2 | GLY | GGC | 27.0 | PRO | CCA | 15.6 |
| ALA | GCC | 11.7 | GLY | GGA | 4.69 | PRO | CCG | 15.6 |
| ALA | GCA | 7.82 | GLY | GGG | 3.13 | SER | UCU | 22.4 |
| ALA | GCG | 7.82 | HIS | CAU | 8.43 | SER | UCC | 14.4 |
| ARG | CGU | 4.21 | HIS | CAC | 13.8 | SER | UCA | 7.28 |
| ARG | CGC | 2.70 | ILE | AUU | 20.3 | SER | UCG | 1.56 |
| ARG | CGA | 2.70 | ILE | AUC | 13.0 | SER | AGU | 3.82 |
| ARG | CGG | 1.56 | ILE | AUA | 3.13 | SER | AGC | 6.26 |
| ARG | AGA | 17.2 | LEU | UUA | 10.9 | THR | ACU | 17.4 |
| ARG | AGG | 1.72 | LEU | UUG | 19.2 | THR | ACC | 11.2 |
| ASN | AAU | 9.56 | LEU | CUU | 0.956 | THR | ACA | 6.26 |
| ASN | AAC | 15.6 | LEU | CUC | 1.56 | THR | ACG | 1.56 |
| ASP | GAU | 15.5 | LEU | CUA | 9.00 | TRP | UGG | 11.7 |
| ASP | GAC | 25.3 | LEU | CUG | 9.00 | TYR | UAU | 10.4 |
| CYS | UGU | 4.56 | LYS | AAA | 6.70 | TYR | UAC | 17.0 |
| CYS | UGC | 7.47 | LYS | AAG | 15.1 | VAL | GUU | 19.3 |
| GLN | CAA | 14.1 | MET | AUG | 7.82 | VAL | GUC | 12.4 |
| GLN | CAG | 1.56 | PHE | UUU | 8.9 | VAL | GUA | 3.13 |
| GLN | GAA | 18.4 | PHE | UUC | 14.6 | VAL | GUG | 2.87 |
| GLN | GAG | 3.13 | PRO | CCU | 3.13 |  |  |  |
| GLY | GGU | 16.5 | PRO | CCC | 2.01 |  |  |  |

Having established that Gilchrist & Wagner’s [47] codon-specific elongation rates are realistic, we used them to simulate the entire budding yeast translatome. The results of our simulations at 99.9% reinitiation are strongly correlated (Fig. 6B) with experimental measures of ribosome densities (r = 0.932, n = 5542, using data from [6]) and calculated initiation rates (r = 0.742, n = 5348, using estimates from [3]; r = 0.618, n = 3728, using estimates from [4]). Our yield predictions are less strongly correlated with measured protein abundances (r = 0.478, n = 4686, data from Peptide Atlas 2013). This weaker correlation is unsurprising as our predictions of yield omit many important determinants of protein abundance including transcript abundance and protein stability. Results of simulations at other reinitiation levels are included in Fig. S3 (fixed transcript lifetime) and Fig. S4 (experimentally measured transcript lifetimes).

## Discussion

We have shown that a fixed transcriptome-wide level of ribosome reinitiation can generate both length-dependent translation and a powerful transcript-specific role for codon usage, but only when reinitiation is extremely efficient. The level of reinitiation in live cells is unknown, but multiple studies have established that reinitiation is much more frequent than *de novo* initiation in cell-free systems. Furthermore, if reinitiation benefits the cell, we would expect it to evolve to become highly efficient. Maintaining a large pool of ribosomes consumes a substantial part of a cell's energy budget and selection will favor mechanisms that allow ribosomes to be used efficiently [49]. If, as reported by Nelson & Winkler [30] and Kopeina et al [31], reinitiation of post-termination ribosomes is faster than *de novo* initiation from the free ribosome pool, then efficient reinitiation reduces the amount of "dead time" ribosomes spend in the pool waiting to be recruited [34,50]. Each ribosome in a cell is therefore able to complete more rounds of translation in a given time interval under high levels of reinitiation compared to low levels of reinitiation. Reinitiation levels should be very closely associated with fitness: the translation initiation rate is thought to be the principal determinant of the rate of cell division [51,52]. Consequently, if reinitiation does occur in living cells, it is hard to imagine why it would not work very efficiently. Direct measurement of the level of reinitiation *in* vivo may soon be possible thanks to recent technological advancements enabling selective labeling of ribosomes [53] and the visualization of translation on individual mRNAs [54,55].

A single fixed level of reinitiation is not necessary to explain length-dependent translation; efficient reinitiation is only required on short transcripts (Fig. S5). Studies in living cells have shown that some transcripts are more likely to be associated with translation factors required to form the closed-loop complex than others [56]. If the closed-loop complex is required for efficient reinitiation, then reinitiation levels likely vary between transcripts. More specifically, shorter transcripts likely experience higher levels of reinitiation since they are both more likely to be enriched with closed-loop factors [15,57) and to form more stable closed-loop complexes [58]. Additionally, cellular depletion of both the closed loop factor eIF4G and the translational regulator Asc1/RACK1 has also been shown to have a greater effect on the translation of short transcripts than on long transcripts [13,15]. Using length-dependent reinitiation levels in our simulations allows the empirical relationship between CDS length and ribosome density, effective initiation rate, and protein yield to be captured at an average reinitiation level orders of magnitude smaller (~90%; Fig. S5) than does a fixed reinitiation level (99.9%; Fig. S5).

Beyond acting on global mechanisms, natural selection also operates to maximize the protein yield of transcripts encoded by individual genes (translational efficiency [46]). Selection for increased translational efficiency can not only increase the abundance of a given protein in a cell, but can also maintain protein levels while minimizing the cost of transcription, which has been shown to be an important determinant of fitness in yeast [59]. The strength of selection depends on the magnitude of the effect of a given mutation on translational efficiency; mutations with larger effects are subject to stronger selection. We have shown that the magnitude of the effect on translational efficiency of altering a given parameter by an equal amount can vary with the length of the altered transcript species. Thus, the strength of selection on mutations that affect a given parameter can be length-dependent [46]. For instance, doubling the reinitiation rate of a single transcript species results in a bigger increase in translational efficiency for shorter transcripts (Fig. 3). Mutations affecting the reinitiation rate of short transcripts are therefore more likely to be selected than are those than occur on long transcripts, potentially contributing to higher levels of reinitiation on shorter transcripts as discussed above (Fig. S5). In contrast, doubling the *de novo* initiation rate does not result in higher translational efficiency on shorter transcripts and, under reinitiation, can actually have smaller effects on shorter transcripts due to increased initiation interference (Fig. 3). Selection for increased translational efficiency on individual transcript species is therefore not predicted to result in higher *de novo* initiation rates on shorter transcripts. Instead, selection under reinitiation will be more effective at reducing initiation interference on shorter transcripts.

At high levels of reinitiation, we have shown that a single slow step in translation causes a greater reduction in the translational efficiency of short transcripts than that of long transcripts (Fig. 5). Eliminating slow steps has larger effects on the translation of short transcripts compared to long transcripts and therefore selection to eliminate slow steps will be most effective in genes encoding short transcripts. Length-dependent selection against slow steps under reinitiation therefore offers an explanation for the negative correlation between codon adaptation and CDS length observed across eukaryotes ([4, 46, 60–64] but see also [65]). Translational efficiency is particularly sensitive to slow sites near the start codon (Fig. 5, see also [21]): slow clearance of the initiation site delays reinitiation (promoting ribosome loss) and blocks *de novo* initiation resulting in lower ribosome densities on affected transcripts. Multiple mechanisms can determine how slowly ribosomes vacate the initiation site including the presence of one or more slow codons [21] or the presence of table 5′ secondary structures in the transcript [66]. As both features reduce yield to a greater extent on short transcripts compared to long transcripts (Fig. 5), selection should be more efficient at eliminating them on shorter transcripts, consistent with the negative correlations between CDS length and both 5′ mRNA folding energy and 5' codon adaptation [60]. Thus, length-dependent translation generated by high levels of reinitiation will generate length-dependent selection against slow steps [46], which will in turn reinforce patterns of length-dependent translation.

Reinitiation provides a simple mechanistic explanation for empirically observed patterns of length-dependent translation including negative correlations between CDS length and ribosome density, effective initiation rate, protein yield, transcript codon adaptation, 5' codon adaptation, 5ʖ folding energy, and association with closed-loop factors. Under reinitiation, these patterns are expected to emerge through selection for efficient ribosome usage, maximizing protein yield, and translational efficiency on individual transcript species. This is in sharp contrast to linear models in which, at low ribosome availability, length-dependence arises through direct selection for higher *de novo* initiation rates on shorter transcripts [3,4]. Our model is consistent with the emerging view that translation is controlled not only by initiation, but also by elongation and termination/reinitiation [21,22,67]. This conceptual shift makes clear that manipulating any these stages can have profound consequences on translation, and presents factors associated with elongation, release, and recycling as new targets for therapeutic intervention (cf. [68]).

## Parameter Estimates and Justification

### Transcript lifetime, age distribution and mode of decay

The decay of eukaryotic transcripts is initiated by the stepwise removal of adenine residues from the poly(A) tail [69]. Deadenylation is the slowest step in mRNA decay, and during this process there is no apparent decay of the transcribed portion of the mRNA [70,71]. Enzymatic digestion of the poly(A) tail is distributive: the deadenylase binds to a poly(A) tail, cleaves a small number of residues, then dissociates from the transcript before binding to a different poly(A) tail [72]. Thus deadenylation consists of a series of sequential first-order reactions, resulting in a hypo-exponential distribution of full-length transcript lifetimes [69,73]. Hypo-exponential distributions, by definition, have lower variances than an exponential function with the same mean lifetime and age-matched populations of eukaryotic transcripts are expected to show very little degradation for long periods then decay rapidly. When the number of sequential deadenylation steps is high (~30), the distribution of transcript lifetimes becomes symmetrical and approximates the normal distribution [74]. We have therefore, for computational simplicity, assumed that all transcripts in a given simulation have the same fixed lifetime.

In our model, all translation ceases as soon as the transcript lifetime is reached; ribosomes that have a new round of translation are not allowed to finish and therefore do not contribute to the protein yield. Consequently, protein yield in our model does not exactly match the effective initiation rate. There is evidence that eukaryotic mRNA decay is co-translational: transcript degradation in the 5′-3′ direction follows the last translating ribosome allowing all ribosomes to complete their final round [75]. If this is the sole mechanism of eukaryotic transcript decay, then the true yield becomes the effective initiation rate multiplied by the transcript lifetime (a constant in our model).

### *De novo* initiation rates

The average number of ribosomes per transcript is remarkably constant across eukaryotes. Polysome profiling studies estimate that the median *S. cerevisiae* transcript contains 6.0 ribosomes (based on weighted averages for all measured transcripts [6]; an identical average was calculated in [7]) while the average human embryonic kidney HEK293T cell contains 5.6-6.1 ribosomes [11]. Direct counts of the numbers of ribosomes in polysomes [14] using atomic force microscopy provide similar values (*S. cerevisiae* = 6.5 ribosomes per transcript, HEK293T = 8.7 ribosomes per transcript, human MCF-7 cells = 8.3 ribosomes per transcript). Direct polysome counts are almost certainly over-estimates of the average number of ribosomes per transcript as they ignore unoccupied transcripts (approximately 15-30% of mRNAs [6,11]) and mRNAs occupied by a single ribosome. To be consistent with these counts, in all of our models, for any transcript lifetime and reinitiation probability, we have adjusted the *de novo* initiation rate such that the average transcript (CDS length = 400 codons) carries 6 ribosomes (white line on each heatmap in Fig. S2). This causes the *denovo* initiation rate to decrease with increasing reinitiation level and decreasing transcript lifetimes.

Maintaining six ribosomes on a 400-codon-long transcript for transcripts with different lifetimes requires much larger changes to the *de novo* initiation rate at high reinitiation probabilities compared to low reinitiation levels (Fig. S2). Under perfect reinitiation, the lifetime protein yields of transcripts of a given CDS length will be similar, but the rate of protein production over time (proteins per mRNA per unit time) will be higher for short-lived transcripts compared to long-lived transcripts. Thus, organisms with short-lived transcripts, such as yeast, will exhibit much higher protein synthesis rates than organisms with long-lived transcripts, such as mammals. Current estimates suggest that protein production rates in mammals are considerably lower than in yeast, consistent with the much higher doubling rates and much lower protein stabilities in yeast compared to mammals [76–78]. In contrast, assuming similar elongation rates across species, linear models predict that yeast and mammals should have similar protein production rates since they have similar average ribosome densities.

### Model parameters

#### Full model

We explored the consequences of different reinitiation levels on the average ribosome density, effective initiation rate and protein yield for transcripts with different CDS lengths using a wide range of transcript lifetimes and *de novo* initiation rates (Fig. S2). For each combination of transcript lifetime and *de novo* initiation rate, we simulated translation for transcripts of the following log-uniform distributed CDS lengths (in codons): 50, 63, 79, 100, 126, 158, 200, 251, 316, 399, 502, 632, 796, 1002, 1263, 1590, 2002, 2522, 3176, and 4000, and then calculated the slope of the resulting estimates of ribosome density, effective initiation rate, and protein yield over CDS length. All codons were decoded at a rate of 10s^-1^ based on the average level in yeast [79] and similar to the average rate observed in a mouse embryonic cell line (5.6s^-1^ [80]). Termination rates (the sum of the release rate and the reinitiation rate) are set equal to the elongation rate at 10s^-1^. Different reinitiation levels are achieved by setting the reinitiation rate as the corresponding proportion of the termination rate. Translation of each transcript was averaged over 1000 runs.

#### General model

To explore the consequences of different reinitiation levels in more detail, we present a model using an arbitrary lifetime of 3000s (50 minutes) for all transcripts (Figs. 2-5). This value is intermediate between estimates of median transcript half-lives in yeast (10-30 minutes [81]) and mammalian cell lines (300-600 minutes [81]). Simulations were performed with a constant elongation rate of 10s^-1^ (except for Fig. S3 where we perform the same simulation at 5s^-1^ and 20s^-1^). The model is otherwise the same as the full model, except that a single *de novo* initiation rate is used at each reinitiation level. *De novo* initiation rates are adjusted for each reinitiation level such that a 400-codon-long transcript carries an average of 6 ribosomes. The *de novo* initiation rates used with a transcriptome-wide elongation rate of 10s^-1^ were: 100% = 0.00438s^-1^, 99.9% = 0.00458s^-1^, 99% = 0.00586s^-1^, 95% = 0.01289s^-1^, 90% = 0.02285s^-1^, 80% = 0.04199s^-1^, 50% = 0.09570s^-1^, 0% = 0.17578s^-1^.

#### Yeast-specific model

We computed the average ribosome density, effective initiation rate, and protein yield (Fig. 6) for 5888 *S. cerevisiae* transcripts ranging in CDS length from 16 to 4910 codons (median length = 405 codons) in our model. We used the codon-specific elongation rates calculated by Gilchrist & Wagner [47]; these rates are scaled such that the average elongation rate is 10s^-1^. As above, we adjusted *de novo* initiation rates for each reinitiation level such that a 400-codon-long transcript (ignoring variation in decoding rates) contained an average of 6 ribosomes. The exact *de novo* initiation rates used were: 100% = 0.00859s^-1^, 99.9% = 0.00869s^-1^, 99% = 0.01016s^-1^, 95% = 0.01641s^-1^, 90% = 0.02568s^-1^, 80% = 0.04492s^-1^, 50% = 0.09766s^-1^, 0% = 0.17578s^-1^.

Most studies of mRNA stability report transcript half-lives. If eukaryotic transcripts decay with biphasic (slow-then-fast) kinetics, then transcript lifetimes cannot be calculated from observed half-lives by assuming first-order kinetics [71]. We have therefore based our estimate of transcript lifetime on a study of nascent transcription rates in *S. cerevisiae* which estimated that the entire set of mRNAs in a cell turns over more than four times per 6780s (113 minute) cell cycle [39], resulting in an average transcript lifetime of 1553s (26 minutes). Although most yeast studies predict fairly similar median transcript half-lives, gene-specific estimates show little correlation across studies [81]. Consequently, we have made the simplifying assumption that all transcripts have the same 1553s lifetime.

## Supporting Information

**Figure S1.**
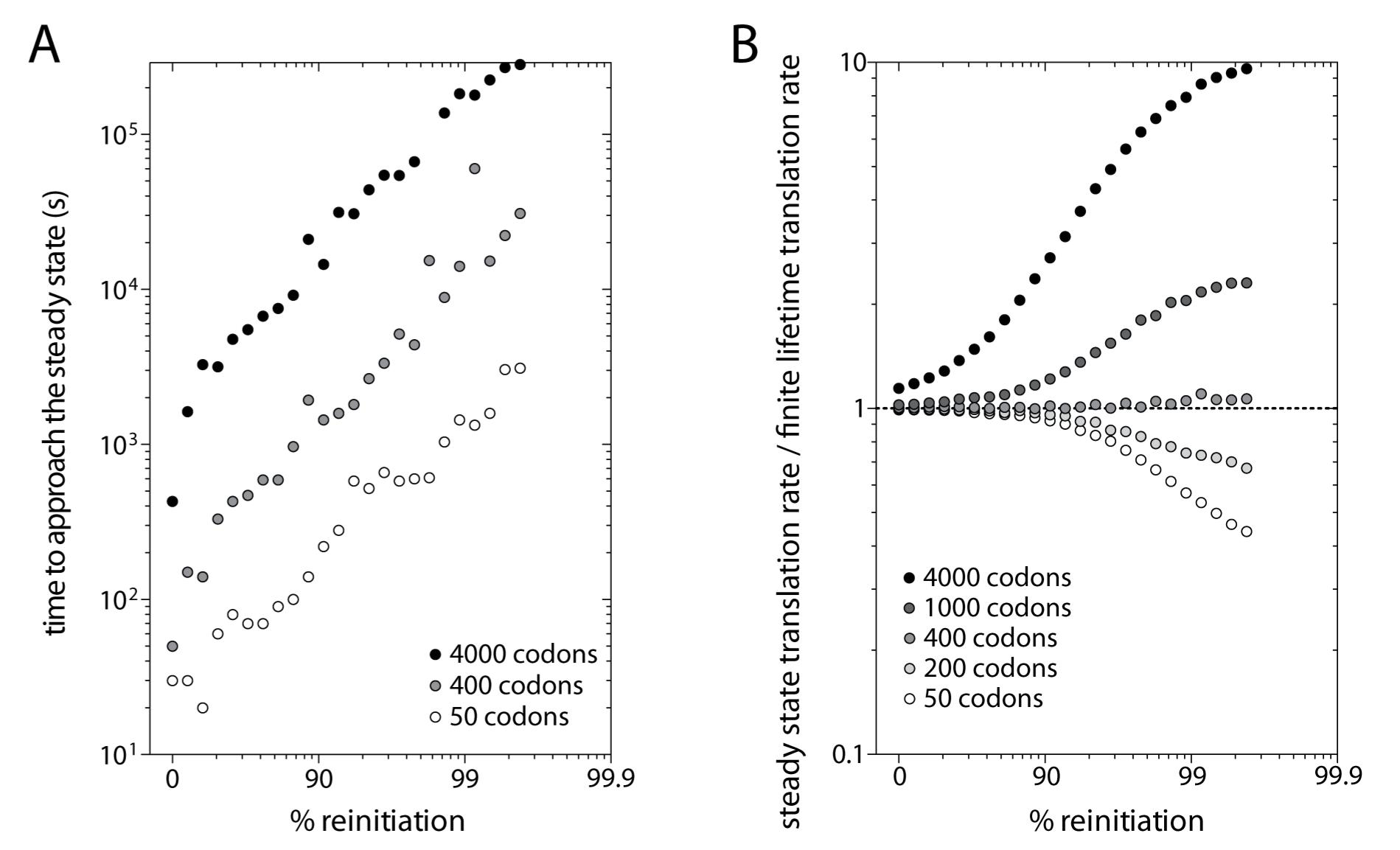
The steady state is a poor approximation of translation at high reinitiation levels for transcripts with finite lifetimes. (A) Time to approach the steady state ribosome density at different reinitiation levels for transcripts of different lengths. Simulations were run for 3×10^5^ s using the parameter values described for our general model. *De novo* initiation rates were adjusted at each reinitiation level so that a 400-codon transcript carried an average of 6 ribosomes at the steady state. The steady state ribosome density was calculated as the average density over the final 10^4^ s of 1000 iterations (if this value showed no directional change over time). The first passage time represents the earliest time point that the average density of 1000 iterations equaled or exceeded the steady state density. Long transcripts (4000 codons) failed to reach the steady state during the 3x10^5^ s run time at reinitiation levels above 99.6%. Data points at higher reinitiation levels were therefore excluded. (B) Steady state translation rate (yield s^-1^) relative to the average translation rate (yield s^-1^) on equivalent transcripts with a finite 3000 s lifetime. Deviations from a ratio of 1 represent the magnitude of the misestimation of the translation rate caused by assuming a perpetual steady state on transcripts with finite lifetimes. *De novo* initiation rates were adjusted at each reinitiation level such that the average 400 codon-transcript carried 6 ribosomes either at the steady state or on average over its 3000 s lifetime. Consequently, all 400 codon transcripts have the same average ribosome density allowing fair comparison. At high reinitiation levels, the steady state approximation overestimates ribosome density on long transcripts and underestimates ribosome density on short transcripts.

**Figure S2.**
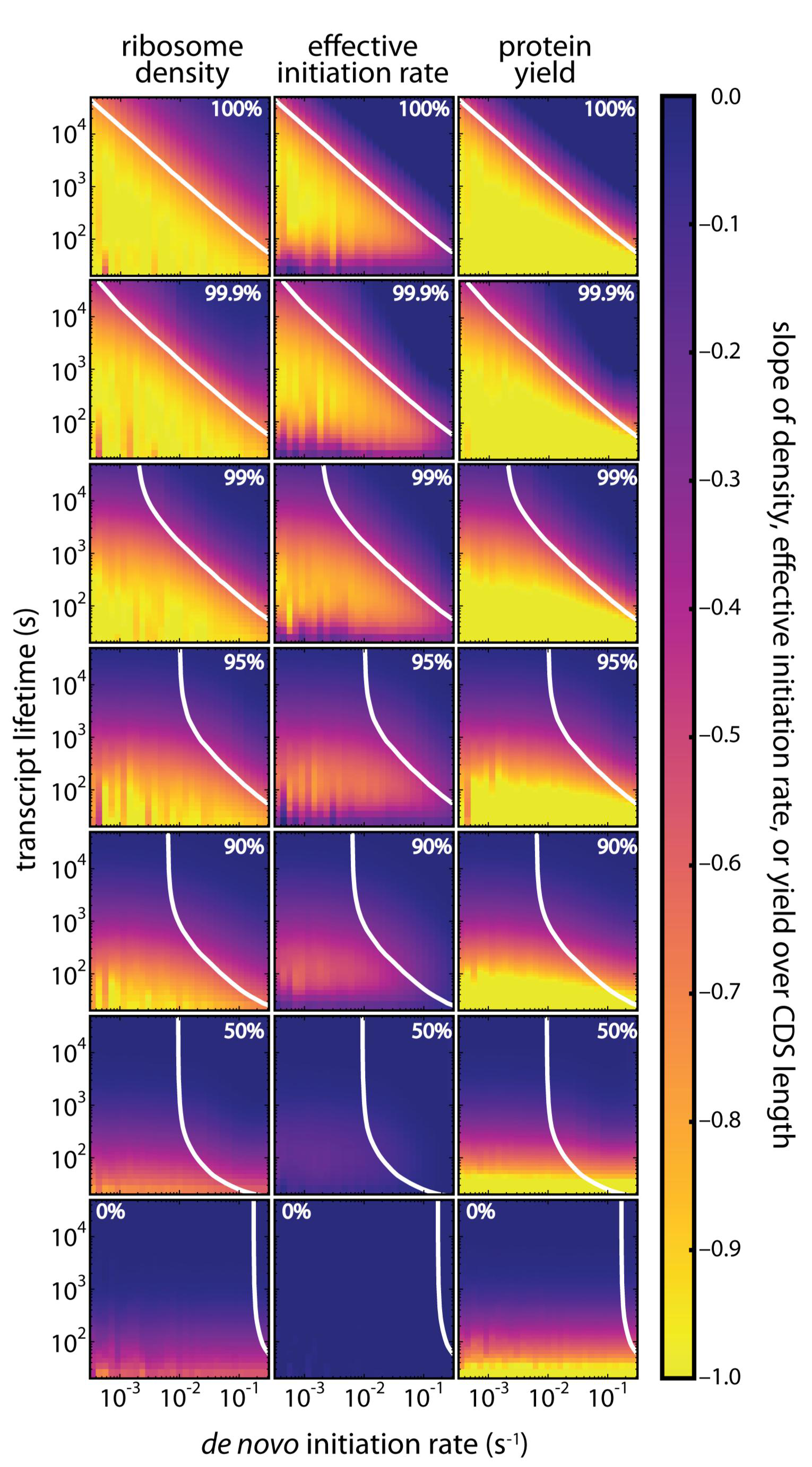
Full model predictions of ribosome density, effective initiation rate, and protein yield at different reinitiation levels. We simulated translation for a wide range of transcript lifetimes and *de novo* initiation rates. For any given combination of transcript lifetime and *de novo* initiation rate, we simulated translation for transcripts with different CDS lengths and then calculated the slope of ribosome density, effective initiation rate, and protein yield over CDS length. Slopes are indicated in different colours (see colour bar), reflecting the degree of length-dependence. The white line in each panel shows the *de novo* initiation rate required at each lifetime such that a 400-codon long transcript carries 6 ribosomes. Our model predicts that high levels of reinitiation can cause length-dependent translation across a wide range of transcript lifetimes and *denovo* initiation rates. At high levels of reinitiation, length-dependence is strongest for short transcript lifetimes andlow *de novo* initiation rates; since ribosomes are unlikely to leave the transcript, at long lifetimes or high *de novo* initiation rates, all transcripts eventually become saturated with ribosomes. As a result, the number of ribosomes per transcript becomes proportional to CDS length (since longer transcripts can carry more ribosomes) and ribosome density, the effective initiation rate, and protein yield become similar for all CDS lengths. As the reinitiation level falls, the effective initiation rate becomes dominated by the *de novo* initiation rate, which is the same for all CDS lengths, and ribosome density, the effective initiation rate, and protein yield all become independent of CDS length. In a linear model of translation (0% reinitiation), negative slopes for density and yield are only seen for extremely short transcript lifetimes simply because the first initiating ribosome fails to reach the stop codon on long transcripts before the maximum lifetime is reached. This effect disappears when transcript lifetimes exceed the time required for the first ribosome to reach the stop codon of the longest transcript (~400 seconds).

**Figure S3.**
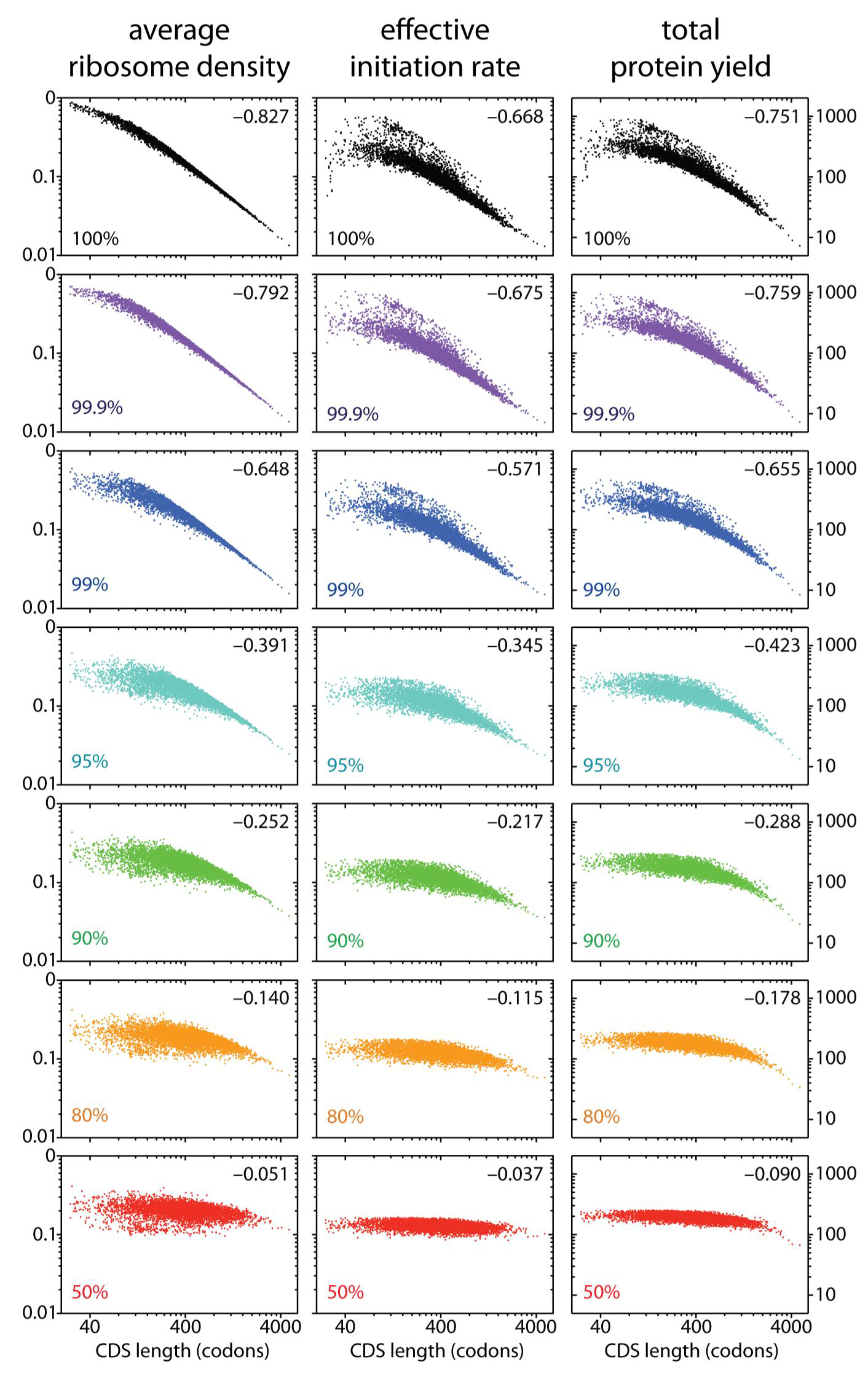
Simulating translation in the budding yeast S. cerevisiae at different reinitiation levels. Predicted ribosome density, effective initiation rate, and protein yield for all 5888 budding yeast transcripts simulated at different reinitiation levels using codon-specific decoding rates from [47]. The left y-axis scale applies to ribosome density (left) and effective initiation rates (center); the right y-axis scale applies to protein yield (right).Slopes are indicated in the top-right corner. As in Fig. 6, all transcripts had a fixed lifetime of 1553 seconds. De novo initiation rates were adjusted at each reinitiation level so that a 400-codon long transcript with a fixed decoding rate of 10s^-1^ carried an average of 6 ribosomes.

**Figure S4.**
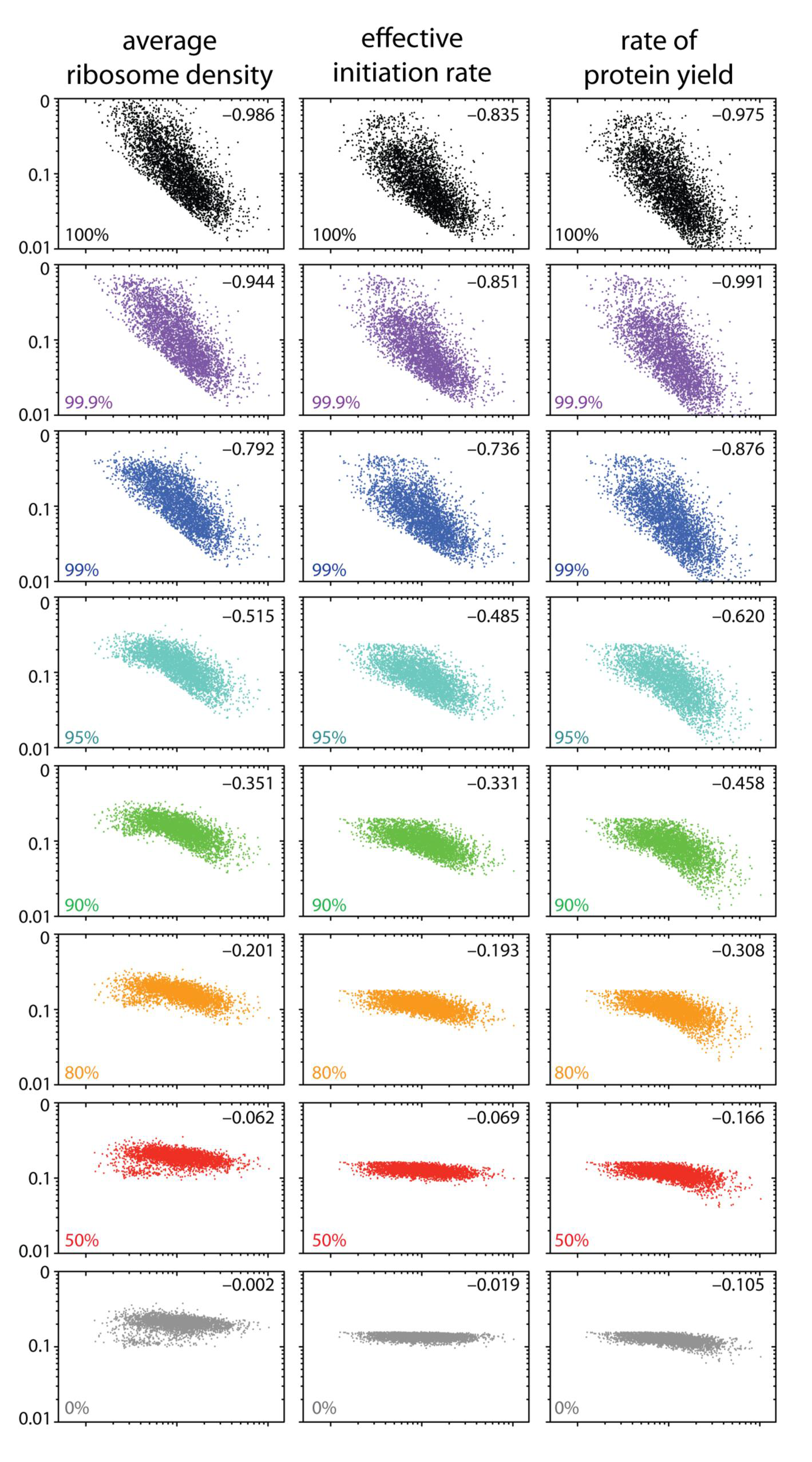
Simulating translation in the budding yeast *S. cerevisiae* at different reinitiation levels using empirical estimates of transcript lifetime. The simulations shown in Fig. S3 were repeated using empirical estimates of transcript lifetimes. Transcript lifetimes were calculated using relative abundances from [82] (measured using single-molecule sequencing digital gene expression which eliminates the length-bias associated with RNA-Seq), a total of 36,000 transcripts per cell [83], and nascent transcription rates from [39]. For each transcript, absolute abundance was divided by the transcription rate to obtain the average lifetime. Transcripts with average lifetimes below 400s were excluded to prevent bias towards increased length-dependence (see Fig. S2).

**Figure S5.**
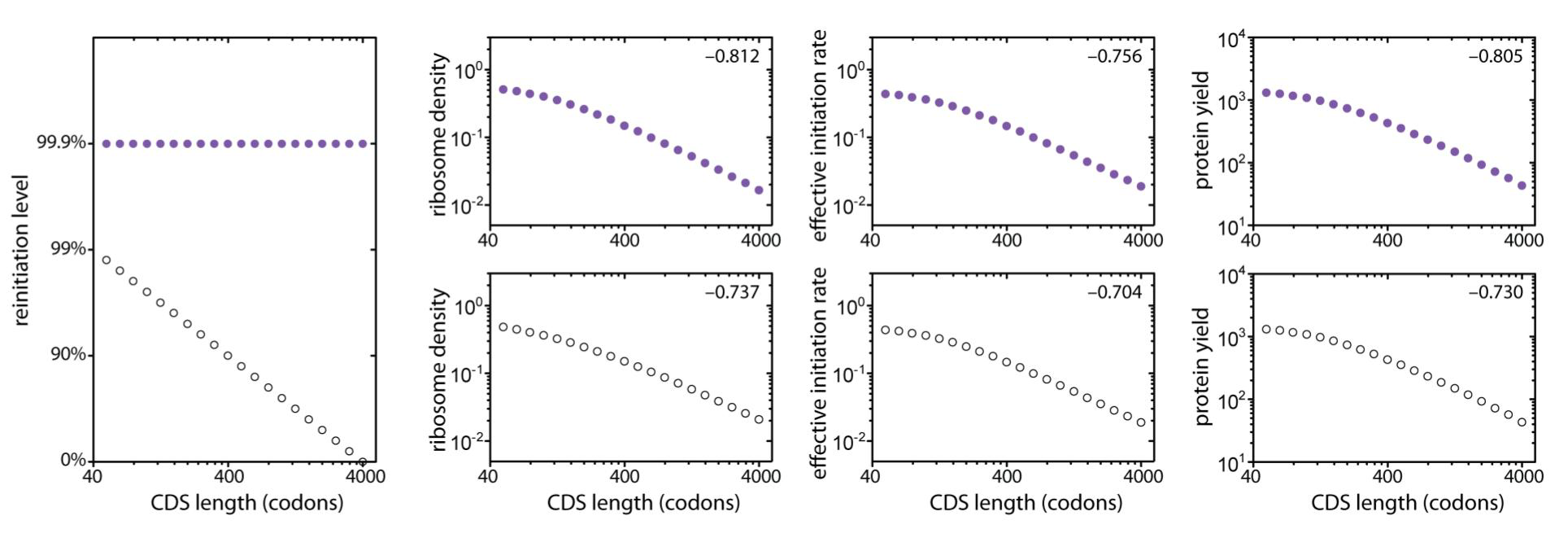
Figure S5. Length-dependent translation only requires high levels of reinitiation on short transcripts. Decreasing reinitiation levels with transcript length (open circles) produces very similar relationships between CDS length and ribosome density, effective initiation rate, and protein yield as does a fixed reinitiation level of 99.9% (purple circles). Slopes of each relationship are shown in the upper right corner of each panel. Length-dependent reinitiation levels capture length dependent translation at much lower reinitiation levels than that required using a fixed reinitiation level. We have arbitrarily chosen to decrease the reinitiation level as a function of CDS length according to the formula 100%*(1- CDS length/4000). The *de novo* initiation rate was set such that 400-codon transcript carried an average of 6 ribosomes (99.9% reinitiation = 0.00458s^-1^, length-dependent reinitiation = 0.02285s^-1^).

